# Pinpoint: trajectory planning for multi-probe electrophysiology and injections in an interactive web-based 3D environment

**DOI:** 10.1101/2023.07.14.548952

**Authors:** Daniel Birman, Kenneth J. Yang, Steven J. West, Bill Karsh, Yoni Browning, the International Brain Laboratory, Joshua H. Siegle, Nicholas A. Steinmetz

## Abstract

Targeting deep brain structures during electrophysiology and injections requires intensive training and expertise. Even with experience, researchers often can’t be certain that a probe is placed precisely in a target location and this complexity scales with the number of simultaneous probes used in an experiment. Here, we present *Pinpoint*, open-source software that allows for interactive exploration of stereotaxic insertion plans. Once an insertion plan is created, Pinpoint allows users to save these online and share them with collaborators. 3D modeling tools allow users to explore their insertions alongside rig and implant hardware and ensure plans are physically possible. Probes in Pinpoint can be linked to electronic micro-manipulators allowing real-time visualization of current brain region targets alongside neural data. In addition, Pinpoint can control manipulators to automate and parallelize the insertion process. Compared to previously available software, Pinpoint’s easy access through web browsers, extensive features, and real-time experiment integration enable more efficient and reproducible recordings.

## Introduction

The availability of high-density low-cost probes, such as Neuropixels (Jun et al., 2017; Steinmetz et al., 2021), has led to a step change in the scale (Allen et al., 2019; Durand et al., 2022; International Brain Laboratory et al., 2023; Steinmetz et al., 2019; Stringer et al., 2019) and ambition (Abbott et al., 2017; Koch et al., 2022) of modern neuroscience. This significant advance in experimental capability is possible in part due to the experimental support provided by advanced software interfaces which support data acquisition (Karsh, 2016; Siegle et al., 2017) and experimental planning (Fuglstad et al., 2023; Peters, 2022). Here, we introduce Pinpoint, software that enables high-quality large-scale electrophysiology research by rendering anatomical brain atlases, calculating the paths in 3D space for dozens of multi-shank probes, preventing collisions, and interfacing with external software and hardware, all in an intuitive and accessible package. Pinpoint lowers the barrier to experimental planning and disconnects the need for expert anatomical knowledge of the brain from the technical skill required to perform complex simultaneous multi-probe recordings. Pinpoint can be run in a web browser with no installation requirements and has a minimal learning curve, making it useful not only for experiment planning, but also for exploring mouse brain anatomy and for teaching new researchers about how *in vivo* experiments are performed.

## Results

Pinpoint provides users with an interactive 3D scene in which electrophysiology trajectories can be explored within the anatomical context of the mouse brain (Fig. 1). At the center of the 3D scene we render transparent 3D meshes for the major brain structures in the mouse Common Coordinate Framework (CCF) (Wang et al., 2020). 3D models of Neuropixels probes (Jun et al., 2017; Steinmetz et al., 2021) or injection needles can be added to the scene and moved through intuitive click+drag or keyboard interaction. Pinpoint provides visualizations of the brain areas that each probe is passing through via a “channel map” (Fig. 1a) and an “in-plane slice” (top right, Fig. 1b). As output, Pinpoint provides the stereotaxic coordinates needed to perform an insertion (Fig. 1e).

**Figure 1:**
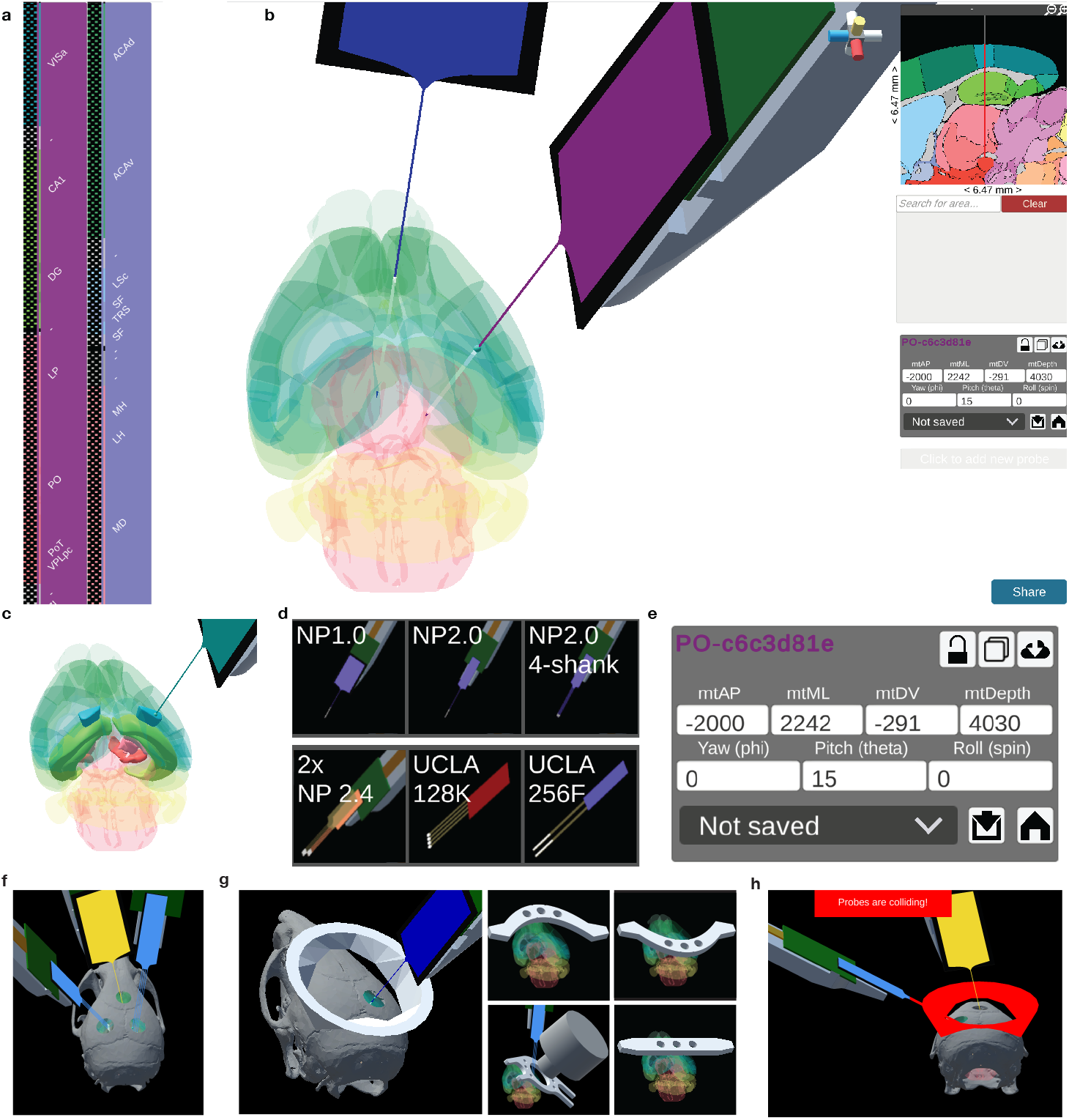
Overview of the Pinpoint interface. (a) Vertical panels show the channel map for two Neuropixels 1.0 probes. Channels are colored by the mouse Common Coordinate Framework region they are inserted in. Area acronyms are shown on the right. (b) The 3D scene shows a transparent view of the mouse brain and models of the Neuropixels probes. On the top right of the user interface (UI), an orientation widget (red, yellow, and blue crosshairs) helps users track the orientation and snap the view to axial, coronal, or sagittal views. The right UI panels include an in-plane slice of the reference atlas, individual area search, stereotaxic coordinates, and at the bottom right a “Share” button which creates a permanent URL to re-load the same scene in another browser. (c) Searching for regions highlights them in the 3D scene as opaque 3D models (highlighted here are parts of visual cortex, hippocampus, and thalamus). (d) Examples of the probes available in Pinpoint. (e) The stereotaxic coordinates relative to a reference coordinate (Bregma, by default) for performing the insertion. The angles (yaw, pitch, and roll) are used to set up the manipulator prior to an experiment. (f) A craniotomy placement tool, combined with a model of the mouse skull, helps users plan surgeries. (g) Example of 3D models that can be placed in the scene, here a “skull cap” used for an implant surgery is shown over the mouse skull, followed by additional examples of experimental hardware including headbars and an imaging lens. (h) Pinpoint detects collisions, to ensure that multi-probe insertion plans are4possible. Colliding models are highlighted in red.

### Planning an insertion trajectory in 3D space

In addition to this brief overview of planning, we also include a video tutorial (Fig. 2) and a comprehensive written tutorial in the Methods. Additional [documentation](https://virtualbrainlab.org/pinpoint/installation_and_use.html) and [in-depth tutorial videos](https://virtualbrainlab.org/pinpoint/tutorial.html) about individual features can be found online.

**Figure 2:**
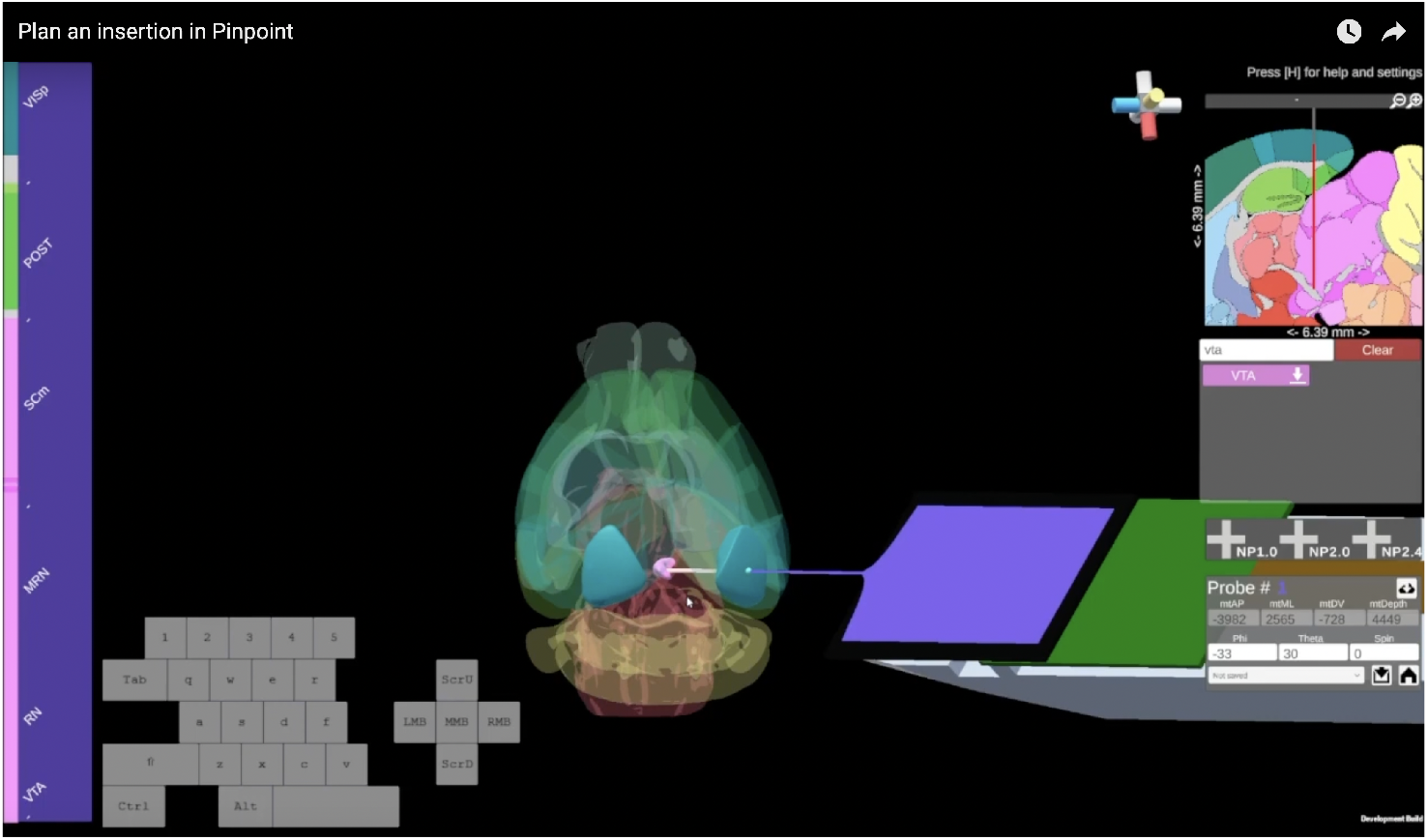
Still frame from the [Plan an insertion in Pinpoint](https://www.youtube.com/watch?v=htTz5tUpCwo) video, a six-minute demonstration of key features.

A central challenge in performing experiments with large-scale electrophysiology is identifying the 3D trajectory required to target the brain region or regions of interest, while constrained by feasible craniotomy locations and experimental apparatus. Visualizing and computing such a trajectory from a 2D paper atlas (Paxinos & Franklin, 2019) is difficult and time-consuming.

To plan an insertion, users need to see both the probe, the areas that they are targeting, and constraints imposed by their setup, while having the ability to optimize probe insertion paths to reach each area of interest. In the default view, Pinpoint displays transparent 3D models of the mouse brain areas defined in the CCF reference atlas. Users can search for areas of interest, highlight these (Fig. 1c), and “snap” probe tips to the center of a particular region. Once a probe is in position, keyboard presses or mouse click+drag controls are used to optimize the trajectory.

To perform the actual implant of a probe, users target the stereotaxic entry coordinate and angles provided by Pinpoint. Pinpoint provides users with the entry coordinate on the brain surface and the depth of the insertion (Fig. 1e) in stereotaxic space. By default, Pinpoint defines the entry coordinate of an insertion relative to Bregma (AP=+5400, ML=+5700, DV=+332 *µ*m in CCF space). Probe angles are defined as yaw (angle around the up axis), pitch (angle off the horizontal plane), and roll (rotation around the probe’s axis).

Because the Allen Common Coordinate Framework (CCF) reference atlas was defined using perfused brains, some users may find that their targeting is improved by using a deformation of this space which better matches the live mouse brain (Fig. 3). In Pinpoint, users can choose to plan their trajectories in the Allen Common Coordinate Framework (CCF, Fig. 3b) space or in a deformation of the CCF (Fig. 3c). The transformed spaces were defined from anatomical MRI images that were aligned to the CCF and in theory provide a better estimate of the live mouse brain, depending on the age and strain. Pinpoint includes two MRI transforms: the Dorr2008 transform (Dorr et al., 2008) taken from the average of 40 in-skull MRIs after death and the Qiu2018 transform (Qiu et al., 2018) taken from the average of 12 live mice. The transforms can be further tailored to individual mice by isometric scaling of the transformed space according to the ratio between an individual mouse’s measured Bregma-Lambda distance relative to the average in the reference atlases, 4150 *µ*m. Additional transforms, such as the Paxinos atlas (Paxinos & Franklin, 2019) will be available in future releases.

**Figure 3:**
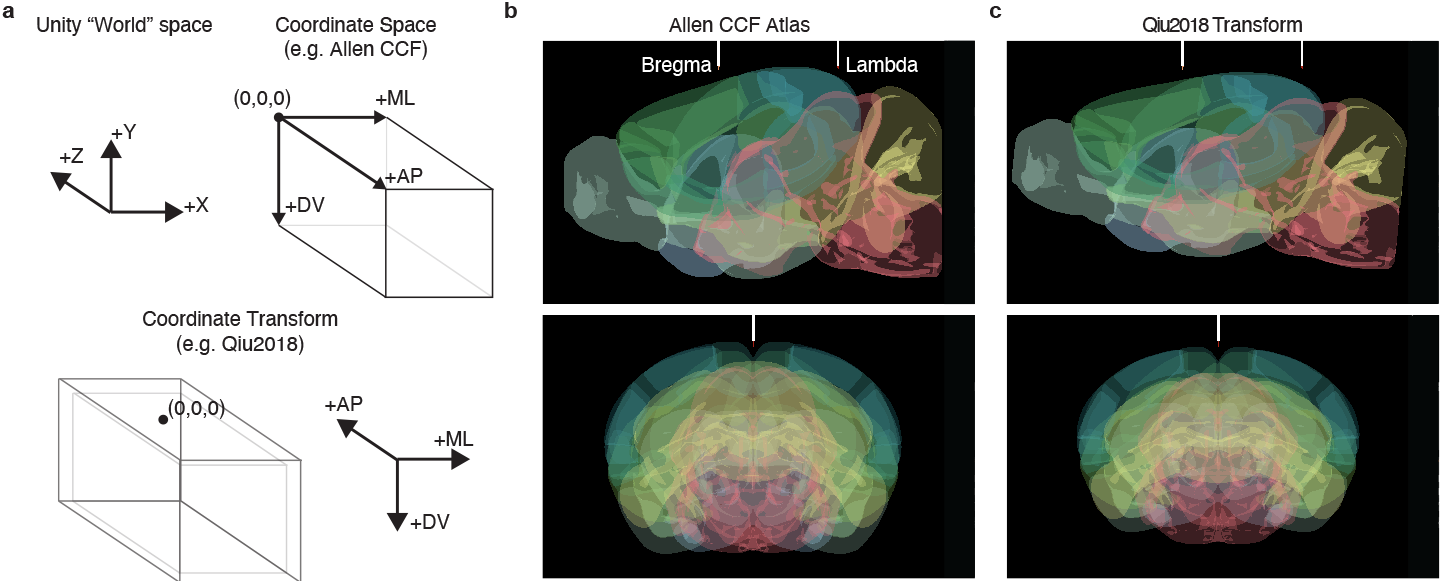
Examples of mouse brain reference atlases included in Pinpoint. (a) Pinpoint supports both histological and anatomically accurate reference atlases. The default axes of the Unity world space represent a standard cartesian coordinate system. To define the space of a reference atlas, such as the Allen Common Coordinate Framework, a Coordinate Space is created, which redefines the zero point to be the top-left-front corner of the reference volume. Because some volumes were not defined using the brains of live animals, we further support Coordinate Transforms, which are affine transforms of an atlas and may optionally redefine the zero coordinate to a different position, such as Bregma, as shown here. (b) Sagittal and coronal slices are shown for the Allen CCF Coordinate Space and (c) for the Qiu et al. (2018) Coordinate Transform, demonstrating how the transform is tilted upwards, stretched along the AP axis and compressed along the DV and ML axes.

When planning complex insertions, users may want to ensure that the 3D geometry of their recording plan will be compatible within the constraints of their craniotomies and experimental hardware. To plan probes relative to craniotomies, Pinpoint can display a skull model and has an interface for adding circular holes of varying radius (Fig. 1f). The 3D scene in Pinpoint can be further modified by adding 3D models for experimental hardware, so long as the position is known relative to Bregma. For example, users can display custom headbars and imaging lenses (Fig. 1g). Pinpoint automatically detects collisions between probes and between probes and hardware and warns the user if a set of insertions will cause problems (Fig. 1h). Adding experimental hardware is not limited to just 3D models, advanced users can use the Unity editor to modify the entire Pinpoint scene to accommodate alternative coordinate systems and rig geometries.

### Interfacing with data acquisition software and hardware

As multi-probe recordings become increasingly complex, there is a need for anatomical targeting information to be available live during recordings. We developed two sets of features in Pinpoint to support this: a hardware application programming interface (API) that allows Pinpoint to communicate with micro-manipulators and a software API that broadcasts the current per-channel annotation data to data acquisition software such as the Open Ephys GUI or SpikeGLX.

We expect users to take advantage of these features in three ways (Fig. 4). First, users can send the planned anatomical targets for an insertion to their data acquisition software as a reference to compare their electrophysiology features against (Fig. 4a). Although this planned anatomical information is not updated as a probe moves, it provides a reference frame that the electrophysiology can be compared against to improve the accuracy of targeting. This can be especially helpful when researchers are targeting regions for the first time, or when performing complex insertions.

**Figure 4:**
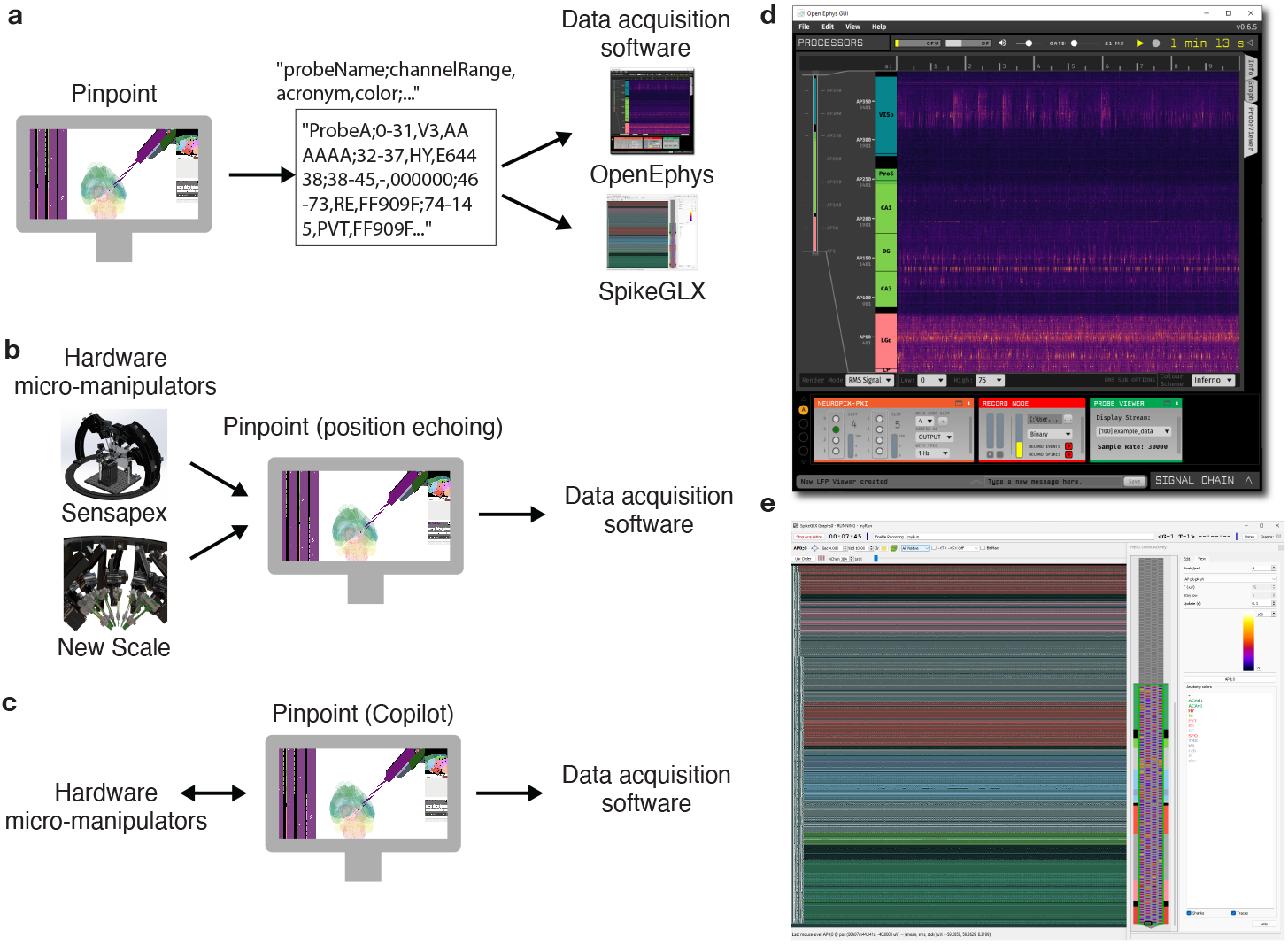
Integration with external hardware and software. Pinpoint exposes two application programming interfaces (APIs) that allow for annotation and position data to be streamed to other applications and for manipulator positions to be echoed in the Pinpoint scene. We present three use cases for these external tools. (a) For researchers who want to improve their targeting accuracy during insertion, Pinpoint can send anatomical information about the final predicted target location of a probe to data acquisition software, such as the Open Ephys GUI or SpikeGLX. Users can compare these per-channel annotations against the electrophysiological signatures they observe on their probe to optimize their targeting. An example of the API channel annotation format is shown in the center box. (b) Pinpoint can also be linked to hardware micro-manipulators from Sensapex and New Scale, showing a live estimate of the current position of the probe during an experiment. This data can then be optionally forwarded to data acquisition software to display a real-time estimate of the probe’s position alongside the electrophysiology. (c) Finally, for users who want to maximize the efficiency of multi-probe recordings and minimize the potential for user error, Pinpoint offers a Copilot mode in which the insertion process is run by the software with minimal user intervention. (d,e) Examples of the Open Ephys and SpikeGLX graphical interfaces displaying probe annotation information.

During a live experiment, researchers often want to know where they are positioning their probe during an insertion. This information helps trainees learn about the brain more efficiently and helps experts avoid missing their target regions. The second way that users can take advantage of the API is by linking a hardware manipulator to Pinpoint and then streaming the probe’s live position to to Open Ephys (Fig. 4d) or SpikeGLX (Fig. 4e) to communicate this information, so that those programs can show the anatomical locations of live electrophysiology data during an experiment. To develop this capability we built a Python package “Electrophysiology Manipulator Link” (Ephys Link) which acts as an intermediate server between Pinpoint and different hardware manipulator platforms. Pinpoint, and other programs, use Ephys Link to read position information from linked manipulators. During an experiment, users place their probe at a reference coordinate (Bregma, by default) and link the live probe to the virtual probe in Pinpoint. Once linked, all movements on the manipulator are reflected in Pinpoint, allowing researchers to see an estimate of their live position in the brain as they insert their probe.

Finally, the hardware API also supports sending commands back to the micro-manipulators, allowing us to control probes both using keyboard commands and to automate the probe insertion process. Pinpoint includes an automation sequence we call “Copilot” that simplifies and parallelizes the process of inserting multiple probes at the same time. Tracking probe positions, logging data about probe movements, and performing the actual movement of probes to their entry coordinates, as well as inserting and removing probes from the brain are all automated, allowing users to focus on other aspects of conducting their experiment. The Copilot sequence does require two manual steps, first when users reference their probes to Bregma, and second when they insert the probe tip into the brain.

### Sharing insertion plans

To plan ongoing complex experiments with many recordings split across multiple days, Pinpoint allows users to save and share their insertion plans. Users can create an account in the Pinpoint system, allowing them to save and load sets of insertions. Users can also at any time create a permanent URL link to a set of insertions, allowing them to share these with collaborators over the internet.

## Discussion

In the past, researchers planned their insertions using 2D slices of the mouse brain, requiring careful calculation to plan complex trajectories across multiple slices. Pinpoint improves on this substantially by allowing users to explore potential probe trajectories through an intuitive 3D interface without any installation requirements. Pinpoint’s hardware and software integration allow users to see their probe position live and to parallelize the insertion process, yielding huge time savings for researchers as they scale up from one or two probes to dozens of simultaneous insertions.

Among Pinpoint’s strengths are its easy access via web browsers, intuitive user interface, and powerful APIs. These advances remain limited by the individual variability of the brain across mice. While Pinpoint can account for isometric scaling that is correlated with skull size through the Bregma-Lambda scaling feature, we are not able to account for nonlinear differences across brains, or uncorrelated variability. Future development could include additional brain atlases, for example developmental mouse atlases (Young et al., 2021) or the Waxholm rat atlas (Papp et al., 2014), and the capability to load individual anatomical MRI data which is critical for targeting in non-human primates.

The future of neuroscience requires that researchers be capable of recording from thousands of neurons, simulta-neously, and across many different model systems. Pinpoint is well-positioned to support this future.

## Methods

### Application and code

Pinpoint is available as a [website](https://data.virtualbrainlab.org/Pinpoint/) or as a standalone desktop application. Desktop releases are distributed through [Steam](https://store.steampowered.com/app/2434260/Pinpoint/). Documentation and tutorials can be found on our [main website](https://virtualbrainlab.org/pinpoint/installation_and_use.html). The code is open-source and can be found on [GitHub](https://github.com/VirtualBrainLab/Pinpoint).

Pinpoint is developed using the Unity Real-Time Development Platform (Unity). Unity is a cross-platform game engine that supports the development of interactive 2D and 3D experiences. We use a number of specific Unity packages to support features in Pinpoint. The base editor (v2021.3.10f1) enables the core features including the 3D scene, use of materials and shaders, and point-and-click and keyboard interaction. The base editor also enables us to distribute Pinpoint for WebGL and the desktop platforms Windows, Linux, and MacOS. The user interface was developed using the Unity UI package. Large assets are bundled and loaded asynchronously using Addressables. Pinpoint also depends on several Unity Asset Store assets: Best HTTP/2, Unisave, Clean Flat UI, and UX Flat Icons.

Some of the functionality developed for Pinpoint is bundled in separate repositories, allowing other users to re-use these specific features in their own projects. The Electrophysiology Manipulator Link server is distributed in the ephys-link repository and releases can be found on [PyPI](https://pypi.org/project/ephys-link/). To support the CCF and other atlases, we developed the [BrainAtlas for Unity](https://github.com/VirtualBrainLab/BrainAtlas) package, which is a wrapper around the BrainGlobe brain-atlas Python package (Claudi et al., 2020).

### Tutorial

The following sections describe the step-by-step process necessary to plan a probe insertion in Pinpoint.

#### Camera controls

Before planning an insertion, it is helpful to master the camera control system. Left click and hold the mouse down in the 3D scene to pitch (Y axis) and roll the brain (X axis). You can also hold space bar while dragging the mouse to yaw the brain. In the top right corner of the scene there is a small axis widget that helps you track the brain’s orientation. The yellow axis is DV (dorsal-ventral), the red axis is AP (anterior-posterior), and the blue axis is ML (medial-lateral, or left-right). Double clicking any of these axes on the widget will snap the brain to the corresponding axial, coronal, or sagittal view, respectively. The scroll wheel on the mouse can be used to zoom in and out. Right click and hold the mouse down to pan the camera and double right-click to reset this.

#### Select a probe

To add a probe to the scene, select the “add new probe” button and then select your Neuropixels, UCLA, or pipette probe style. The Pinpoint scene supports an unlimited number of probes, but only 16 shanks can be shown at the same time on the user interface. If you add more shanks to the scene, the “Show only active probe panels” setting will be enabled, hiding the channel map data for probes that are not actively selected.

#### Setting up targets

Target regions can be highlighted in the 3D scene by searching for them in the search bar (“Search for area…”, right side of UI) and then clicking on the area in the hierarchy. Clicking on a region again removes it from the 3D scene and all highlighted areas can be removed using the red “Clear” button. When there is an active probe in the 3D scene the “Snap” button (downward pointing arrow) can be used to move the probe tip to the center of a particular brain region. Areas can also be highlighted by clicking them in the in-plane slice view (top right of UI).

#### Keyboard and mouse controls

Once a probe is aligned to a target region, the trajectory will need to be optimized. Users can enter changes in the angles and position of the probe through the stereotaxic coordinate panel (right side of UI), or by using the keyboard or mouse. To use the coordinate panel, type the new AP, ML, DV, Depth, Yaw, Pitch, or Roll value into the corresponding box and press enter to apply the change. Note that Pinpoint always reports back the *entry coordinate* and depth of an insertion. This means that if you enter a target coordinate for the tip of an insertion (with a depth of zero) the values you enter will be replaced in the coordinate panel with the entry coordinate and depth when you apply the change.

To use the keyboard, press the key corresponding to the axis and direction: anterior-posterior, W and S; left-right, A and D; dorsal-ventral, E and Q. Keyboard clicks can also rotate probes: yaw, 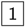 and 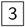 ; pitch, 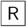 and ; 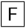 roll, 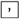 and 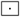. To move the probe along its depth axis press 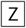 or 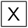. Each of these keyboard presses moves the probe tip by 10 *µ*m along that axis. Holding 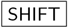 increases the step size to 100 *µ*m m while holding 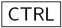 reduces it to 1 *µ*m.

To use the mouse click and drag controls, press and hold the left mouse button on the 3D model of the probe. While holding the mouse button down, press any of the axis keys (e.g. 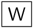 for anterior-posterior). Then, drag the mouse along the axis to move the probe in a continuous motion. Note that the dragging is along the axis, so from some camera views certain axes are not available, for example when looking at the axial plane (from above) it is not possible to drag the probe along the dorsal-ventral axis.

#### Channel map

On the left side of the UI, the channel maps for the active probe will be visible. By default, the channel map uses the bank zero setting for Neuropixels probes or all channels for UCLA probes and pipettes, this can be changed in the Probe settings panel. Channels are colored by the region they are inside of within the CCF hierarchy and the acronyms for each region are shown centered alongside the channels. A thin colored line next to the channel map helps to clearly show the boundaries between regions. Note that the channel map is deformed to match the available UI space and does not correspond to the real-world scaling of the probe.

### Surgery coordinates

Pinpoint computes the entry coordinate for surgeries by finding the coordinate in the reference atlas where the brain surface and the trajectory intersect. Each probe insertion is defined by a tip coordinate (AP, ML, DV) and angles (Yaw, Pitch, Roll). To find the entry coordinate, we step down the probe insertion vector until we are inside the brain. The last coordinate *outside* the brain is considered to be the entry. Raw coordinates are reported in their transformed space, so that they best match the live mouse brain. By default, Pinpoint uses the 25 *µ*m CCF atlas, so entry coordinates will necessarily differ by a small amount from the dura surface but these differences are smaller than the expected variability introduced by experimenters (International Brain Laboratory et al., 2022).

By default, Pinpoint reports the entry coordinate relative to Bregma. This setting can be changed in the Atlas settings, using CCF conventions for the coordinate. For example, to place probes in raw CCF coordinates you would set the reference coordinate to (0,0,0).

To perform a recording, users must level the mouse skull along the Bregma-Lambda and Left-Right axes and set up their probe according to the specified angles. Users then zero their stereotax with the probe tip at the reference coordinate (Bregma, by default) and move the probe to the entry coordinate. If necessary, the probe should be lowered along the DV axis to reach the brain surface. Once at the entry coordinate on the brain surface, users should zero the depth axis and lock all other axes. The target brain area is then reached by driving to the specified depth value. Note that to achieve accurate targeting two references are set: first at Bregma, and then independently at the dura surface for the depth axis.

### Atlases and transformations

We define anatomical atlases in Pinpoint using a CoordinateSpace and a CoordinateTransform. The CoordinateS-pace defines the transformation of coordinates in Unity “World” space into the reference atlas. By default in Unity the Z axis points forward, the X axis to the right, and the Y axis up. The CoordinateSpace for the Allen CCF rotates a vector such that it is redefined according to the anterior-poster, left-right or medial-lateral, and ventral-dorsal axes instead of the world axes. According to the specifications, the CCF is defined such that the (0,0,0) coordinate is at the front, top, left coordinate. By collecting these rotations and offsets into a CoordinateSpace class we make it easy to extend Pinpoint in the future to other anatomical atlases.

Because the CCF was defined from perfused brains it is deformed from the live mouse brain. A CoordinateTrans-form attempts to reverse these deformations and make it possible to examine anatomical data in the space of individual mouse brains. Each CoordinateTransform defines the transformation of a point from a CoordinateSpace into a deformed space defined by anatomical data from a particular mouse or strain, or an average of several mice.

To define each CoordinateTransform we registered the mouse CCF (Wang et al., 2020) atlas into an anatomical MRI space. Pinpoint implements two CoordinateTransforms: the Dorr2008 atlas defined from 40 in-skull anatomical MRIs taken from p84 C57BL/6j mice after death (Dorr et al., 2008) and the Qiu2018 atlas defined from 12 p65 C57BL/6j mice that were alive at the time of anatomical scanning (Qiu et al., 2018). Pinpoint also implements a rotation in each CoordinateTransform to level the Bregma-Lambda surface, pitching the CCF atlas up by 5 degrees (International Brain Laboratory et al., 2022).

Transforms were computed using BrainRegister (West, 2021), a registration pipeline based on elastix (Klein et al., 2010) with optimised parameters for mouse brain registration.

#### Bregma and lambda coordinates

The Bregma and Lambda coordinates in CCF space were determined by finding these points on the average MRI model from Qiu et al. (2018) after alignment to the CCF reference atlas.

### Sharing and Accounts

To save and load experimental plans we developed a feature to share insertion plans through permanent online links and an accounts system that allows users to store insertions in a private database. The share button creates a Base64-encoded query string consisting of the JSON-formatted insertion data for all probes visible in the scene, as well as limited information about the scene view itself.

For the accounts system we integrated Unisave, a back-end database which stores information about insertions in an account secured by a user’s email and a password. Passwords are encrypted according to industry standard practices and are not recoverable. Each account in Unisave is a collection of experiments and each experiment is a collection of insertions and their metadata (e.g. name, probe color).

### Application Programming Interface (API)

Pinpoint exposes an API for sharing per-channel information about each active probe in the scene. The API generates an array of data strings using the format: “[probe0 data, probe 1 data, …]” with each probe’s data formatted using the syntax: “probe-id:channel 0 data;channel 1 data”, and finally with the channel data formatted as “channel#,ccf-atlas-id,ccf-atlas-acronym,hex-color”. The string can be manually exported from the API tab in the settings. Toggling the API features for Open Ephys or SpikeGLX forwards compressed versions of the data string designed for their respective APIs. Each program can then display anatomical information alongisde the electrophysiology data.

### Electrophysiology Manipulator Link (Ephys Link)

To enable position echoing and automated control of micro-manipulators, we developed an intermediate Python package *ephys-link* which handles communication between Pinpoint and various hardware manipulators. Ephys Link is a Python-based WebSocket server split into three layers that connects client applications to manipulator platforms using a standardized set of WebSocket events. The main layer is an asynchronous HTTP server that is responsible for serving the WebSocket connection, error checking, serial connections to peripherals, and managing the user interface. The server calls a manipulator platform layer which defines platform-specific implementations for the declared WebSocket events, manages connected manipulator instances, and handles API calls that affect all instances simultaneously. Each manipulator platform can optionally define a manipulator class that holds details specific to *in vivo* manipulator instances such as its ID, position, axis transform, and movement queue. Creating a client application that uses Ephys Link to communicate with manipulator platforms requires making calls to the defined WebSocket events and handling the returned information.

#### Position echoing

Knowing the position of a probe live during an experiment allows researchers to make substantial changes to their insertion plan while still being confident about reaching their intended brain region target. To support this, *ephys-link* detects and communicates with manipulators that are connected to the same computer and exposes a WebSocket connection for communicating with clients, such as Pinpoint. Once connected, users need to match their probes in the Pinpoint scene to the live manipulators and then set a reference position. To match the probes the yaw, pitch, and roll angles must be set to be identical and the mouse’s skull must be level along the Bregma-Lambda and Left-Right axes. To set the reference, the live probe needs to be moved to the reference coordinate (Bregma, by default) so that Pinpoint can record this position and subtract it from further movements. After linking and referencing, further movements of the micro-manipulator are echoed in the scene allowing users to visualize their probe position live relative to the mouse brain.

#### Copilot

To maximize efficiency and reproducibility of experiments, it’s critical to use an insertion process that is repeatable and which minimizes errors. When connected to the *ephys-link* package, Pinpoint can act as a Copilot during the insertion process to achieve perfect repetition of pre-specified insertion plans at safe insertion speeds. To use Copilot in Pinpoint we developed a standard sequence for performing multi-probe insertions: first, users set up the Pinpoint scene to have the 3D models of all probes. Second, they link these probes to the hardware manipulators and choose the target insertions. Third, they zero all of the probes at Bregma individually. Finally, they allow the insertion sequence to run. The sequence moves probes first to their insertion coordinate, allows the user to enter the brain, and finally drives the probes to the appropriate depth at a speed which minimizes damage during the insertion (Fiáth et al., 2019) and includes time for settling. Note that brain entry step is manual: we don’t support automated entry because probes often do not enter smoothly and require user input to avoid damage to probes. After a recording, probes are automatically retracted in parallel, freeing up researchers to focus on other tasks.

### Other settings

In addition to the features and settings already described, Pinpoint also features a number of other optional settings that make planning insertions more convenient. Users can choose to change the coordinate system to be aligned to the axis of their probe, automatically converting from AP/ML coordinates to their probe’s Forward/Side axes. Pinpoint defines a probe’s angles to be at (0,0,0) when the probe tip is pointing anterior with the electrode surface pointing up, but a setting allows for alternate conventions such as those used by the IBL (where 0,0,0 is a probe pointing ventral with the electrode surface pointing left). Area settings allow users to switch between using area acronyms or full names and to drop white matter tracts that are unlikely to be targeted during an experiment. Atlas settings allow users to switch transforms and to turn on coronal and sagittal slices which move with the probe tip, helping to better align probes within the brain. A coverage atlas (saved as a flattened numpy byte array using np.to_bytes(), where 0 = transparent and 1/2/… indicate increasing coverage) can also be loaded from the Atlas settings from any URL and overlaid onto the reference atlas. Graphics settings allow users to change the background color, make inactive probes transparent, hide the entry coordinate on the brain surface, and hide brain regions that are not being targeted.

### Contributing

Pinpoint is an open-source project and is open to reports about bugs as well as contributions from the neuroscience community. Bugs, comments, and concerns can be reported through the [issues page](https://github.com/VirtualBrainLab/Pinpoint/issues). Instructions for developing and contributing to Pinpoint can be found on our [development page](https://virtualbrainlab.org/pinpoint/development.html).

## Funding

We acknowledge the generous support of the Washington Research Foundation Postdoctoral Fellowship to DB, the Research Scholarship from the Mary Gates Endowment for Students to KY, the Wellcome Trust (216324),

Simons Foundation, and NIH Grant U19NS123716 to the IBL.

## Acknowledgements

We thank Kai Nylund for help designing an early prototype, the Steinmetz Lab and International Brain Laboratory for feedback on the user experience and interface, and Sensapex and New Scale Technologies for their assistance with SDK access to their hardware platforms.

## Contributions

DB -conceptualization, software, supervision, funding acquisition, visualization, methodology, project administration, writing -original draft & review and editing, KY -software, visualization, SJ -formal analysis, methodology, BK -software, JS -software, writing -review and editing, YB -methodology, software, IBL -funding acquisition, writing -review and editing, NS -conceptualization, supervision, funding acquisition, writing -review and editing.

## Notes

https://virtualbrainlab.org/pinpoint/installation_and_use.html

https://github.com/VirtualBrainLab/Pinpoint

### Competing Interest Statement

The authors have declared no competing interest.

